# A large, open source dataset of stroke anatomical brain images and manual lesion segmentations

**DOI:** 10.1101/179614

**Authors:** Sook-Lei Liew, Julia M. Anglin, Nick W. Banks, Matt Sondag, Kaori L. Ito, Hosung Kim, Jennifer Chan, Joyce Ito, Connie Jung, Nima Khoshab, Stephanie Lefebvre, William Nakamura, David Saldana, Allie Schmiesing, Cathy Tran, Danny Vo, Tyler Ard, Panthea Heydari, Bokkyu Kim, Lisa Aziz-Zadeh, Steven C. Cramer, Jingchun Liu, Surjo Soekadar, Jan-Egil Nordvik, Lars T. Westlye, Junping Wang, Carolee Winstein, Chunshui Yu, Lei Ai, Bonhwang Koo, R. Cameron Craddock, Michael Milham, Matthew Lakich, Amy Pienta, Alison Stroud

## Abstract

Stroke is the leading cause of adult disability worldwide, with up to two-thirds of individuals experiencing long-term disabilities. Large-scale neuroimaging studies have shown promise in identifying robust biomarkers (e.g., measures of brain structure) of long-term stroke recovery following rehabilitation. However, analyzing large rehabilitation-related datasets is problematic due to barriers in accurate stroke lesion segmentation. Manually-traced lesions are currently the gold standard for lesion segmentation on T1-weighted MRIs, but are labor intensive and require anatomical expertise. While algorithms have been developed to automate this process, the results often lack accuracy. Newer algorithms that employ machine-learning techniques are promising, yet these require large training datasets to optimize performance. Here we present ATLAS (Anatomical Tracings of Lesions After Stroke), an open-source dataset of 304 T1-weighted MRIs with manually segmented lesions and metadata. This large, diverse dataset can be used to train and test lesion segmentation algorithms and provides a standardized dataset for comparing the performance of different segmentation methods. We hope ATLAS release 1.1 will be a useful resource to assess and improve the accuracy of current lesion segmentation methods.

## Background & Summary

Approximately 795,000 people in the United States suffer from a stroke every year, resulting in nearly 133,000 deaths^1^. In addition, up to 2/3 of stroke survivors experience long-term disabilities that impair their participation in daily activities^2,3^. Careful clinical decision making is thus critical both at the acute stage, where interventions can spare neural tissue or be used to promote early functional recovery^4^, and at the subacute/chronic stages, where effective rehabilitation can promote long-term functional recovery. Enormous efforts have been made to predict outcomes and response to treatments at both acute and subacute/chronic stages using brain imaging.

At the acute stage, within the first 24 hours or so after stroke onset, clinicians face important, time-sensitive decisions such as whether to intervene to save damaged tissue (e.g., administer thrombolytic drugs, perform surgery). Clinical brain images such as magnetic resonance imaging (MRI) and computerized tomography (CT) scans are routinely acquired to help diagnose and make these urgent clinical decisions. Images obtained often include lower-resolution CT scans or structural MRIs (e.g., T2-weighted, FLAIR, diffusion weighted, or perfusion weighted MRIs), and impressive efforts have been made to use these images to automatically detect the lesion volume, predict responses to acute interventions, and predict general prognosis. As clinical scans are typically a mandatory part of acute stroke care, there has been excellent progress in using large-scale datasets of the acquired images to relate to outcomes and build automated lesion detection algorithms and predictive models over the past few decades^5^. In addition, using imaging to assess the extent of neural injury within the first few days after stroke can be helpful for informing entry criteria and stratification variables for enrollment in clinical trials of early recovery therapies, which have specific time windows shortly after stroke onset^4^.

On the other hand, there have been fewer advances in large-scale neuroimaging-based stroke predictions at the subacute and chronic stages. Here, clinicians must triage patients and assign scarce rehabilitation resources to those who are most likely to benefit and recover. Brain imaging, such as MRI, is primarily acquired as part of research studies to understand brain-related changes in response to different therapeutic interventions or to provide valuable additional information, beyond what can be gleaned from bedside exams, that can be used to predict rehabilitation outcomes^6^. As stroke is a leading cause of adult disability worldwide, there is a large emphasis placed on predicting and understanding how to best promote long-term rehabilitation in these individuals. Although there are fewer MRIs acquired during this time, the most common research scan is a high-resolution T1-weighted structural MRI, which is often acquired along with functional MRI and high-resolution diffusion MRI scans and can show infarcts at the post-acute stage. Research using these types of images at this stage of stroke have shown promising biomarkers that could potentially provide additional information, beyond behavioral assessments, to predict an individual’s likelihood of recovery for specific functions (e.g., motor, speech) and response to treatments^7–9^. Thus far, measures that include the size, location, and overlap of the lesion with existing brain regions or structures, such as the corticospinal tract, have been successfully used as predictors of long-term stroke recovery and rehabilitation^9–15^. However, to date, this has only been done in smaller-scale studies, and results may conflict across studies or be limited to each sample. Examining lesion properties with larger datasets at the subacute and chronic stages could lead to the identification of more robust biomarkers for rehabilitation that are widely applicable across diverse populations. Recently, efforts for creating large-scale stroke neuroimaging datasets across all time points since stroke onset have emerged and offer a promising approach to achieve a better understanding of the long-term stroke recovery process (e.g., ENIGMA Stroke Recovery; http://enigma.ini.usc.edu/ongoing/enigma-stroke-recovery/).

However, a key barrier to properly analyzing these large-scale stroke neuroimaging datasets to predict rehabilitation outcomes is accurate lesion segmentation. While many acute neuroimaging stroke studies bypass manual lesion segmentation by using a visual scoring of lesion characteristics with validated scoring tools applied by expert raters, research studies that wish to examine the overlap of the lesion with specific brain structures (e.g., in voxel-based lesion symptom mapping, or lesion load methods) require an accurate and detailed lesion map. In T1-weighted MRIs, which are often used in research, the gold standard for delineating these lesions is manual segmentation, a process that requires skilled tracers and can be prohibitively time consuming and subjective^16^. A single large or complex lesion can take up to several hours for even a skilled tracer. As a result of this demand on time and effort, this method, which has been used in previous smaller neuroimaging studies, is not suitable for larger sample sizes. Based on the literature, most studies with manually segmented brain lesions on T1-weighted MRIs use smaller sample sizes between 10 to just over 100 brains^16–20^. Accurately segmenting hundreds or thousands of stroke lesions from T1-weighted MRIs may thus present a barrier for larger-scale stroke neuroimaging studies.

Many stroke neuroimaging studies have utilized semi- or fully-automated lesion segmentation tools for their analyses. Semi-automated segmentation tools employ a combination of automated algorithms, which detect abnormalities in the MR image, and manual corrections or inputs by an expert. Fully-automated algorithms rely completely on the algorithm for the lesion segmentation. While these require little human input or expertise, they still may require significant computational resources and processing time.

Many of these fully-automated algorithms employ machine learning techniques that require training and testing on large datasets^21^, and the performance of the algorithm is highly dependent on the size and diversity of the training dataset. While there have been several exciting initiatives regarding lesion segmentation in acute clinical imaging, discussed below, there are few publically available large training/test datasets of manually segmented stroke lesion masks on research-grade T1-weighted images that could be used for improving such algorithms. Thus, while both semi- and fully-automated lesion segmentation tools have the potential to greatly reduce the time and expertise needed to analyze stroke MRI data^22^, it is unclear whether they provide the accuracy needed for rigorous stroke lesion-based analyses.

In addition, it is difficult to compare the performance of automated lesion segmentation tools as they are often not evaluated for performance on the same dataset. Recently, some exciting initiatives have emerged to develop better segmentation algorithms using standardized datasets and metrics. In particular, the Ischemic Stroke Lesion Segmentation (ISLES) challenge is an annual satellite challenge of the Medical Image Computing and Computer Assisted Intervention (MICCAI) meeting that provides a standardized multimodal clinical MRI dataset of approximately 50-100 brains with manually segmented lesions^23^. The ISLES competition encourages research groups to use the dataset to evaluate their lesion segmentation algorithms and predict acute outcomes to inform clinical decision making. This approach is promising for developing better lesion segmentation algorithms and predictive models for acute imaging. However, past ISLES challenge datasets have traditionally focused more on using multimodal clinical MRIs to predict more acute results, and these algorithms are not easily translatable to the high-quality T1-weighted MRIs typically found in subacute/chronic stroke rehabilitation research. Thus, here, we aimed to develop a complementary large dataset using only anatomical T1-weighted MRIs, which are typically acquired in research studies after the acute stage to assess rehabilitation outcomes. We anticipate this dataset could be useful for enhancing lesion segmentation methods for T1-weighted images often used in medical rehabilitation research.

Here, we present ATLAS (Anatomical Tracings of Lesions After Stroke) Release 1.1, an open-source dataset consisting of 304 T1-weighted MRIs with manually segmented diverse lesions and metadata. The goal of ATLAS is to provide the research community with a standardized training and testing dataset for lesion segmentation algorithms on T1-weighted MRIs. We note that this dataset is not representative of the full range of stroke, as this data was acquired through research studies in which individuals with stroke voluntarily participated, and all participants had to be eligible for a research MRI session. However, this dataset may be useful for testing and comparing the performance of different lesion segmentation techniques and identifying key barriers hindering the performance of automated lesion segmentation algorithms. We believe that this diverse set of manually segmented lesions will serve as a valuable resource for researchers to use in assessing and improving the accuracy of lesion segmentation tools.

## Methods

### Data Overview

304 MRI images from 11 cohorts worldwide were collected from research groups in the ENIGMA Stroke Recovery Working Group consortium. Images consisted of T1-weighted anatomical MRIs of individuals after stroke. These images were collected primarily for research purposes and are not representative of the overall general stroke population (e.g., only including individuals who opt in to participate in a research study, and excluding individuals with stroke who cannot undergo MRI safely).

For each MRI, brain lesions were identified and masks were manually drawn on each individual brain in native space using MRIcron^24^, an open-source tool for brain imaging visualization and defining volumes of interest (http://people.cas.sc.edu/rorden/mricron/index.html). At least one lesion mask was identified for each individual MRI. If additional, separate (non-contiguous) lesions were identified, they were traced as separate masks. An expert neuroradiologist reviewed all lesions to provide additional qualitative descriptions of the type of stroke, primary lesion location, vascular territory, and intensity of white matter disease. Finally, a separate tracer performed quality control on each lesion mask. This included assessing the accuracy of the lesion segmentations, revising the lesion mask if needed, and categorizing the lesions to generate additional data such as the number of lesions in left and right hemispheres, and in cortical and subcortical regions. This dataset is provided in native subject space and archived (n=304). A subset of this dataset was also defaced, intensity normalized, and provided in standard space (normalized to the MNI-152 template, n=229; for an overview of the dataset and archives, see Figure 1).

**Figure 1.**
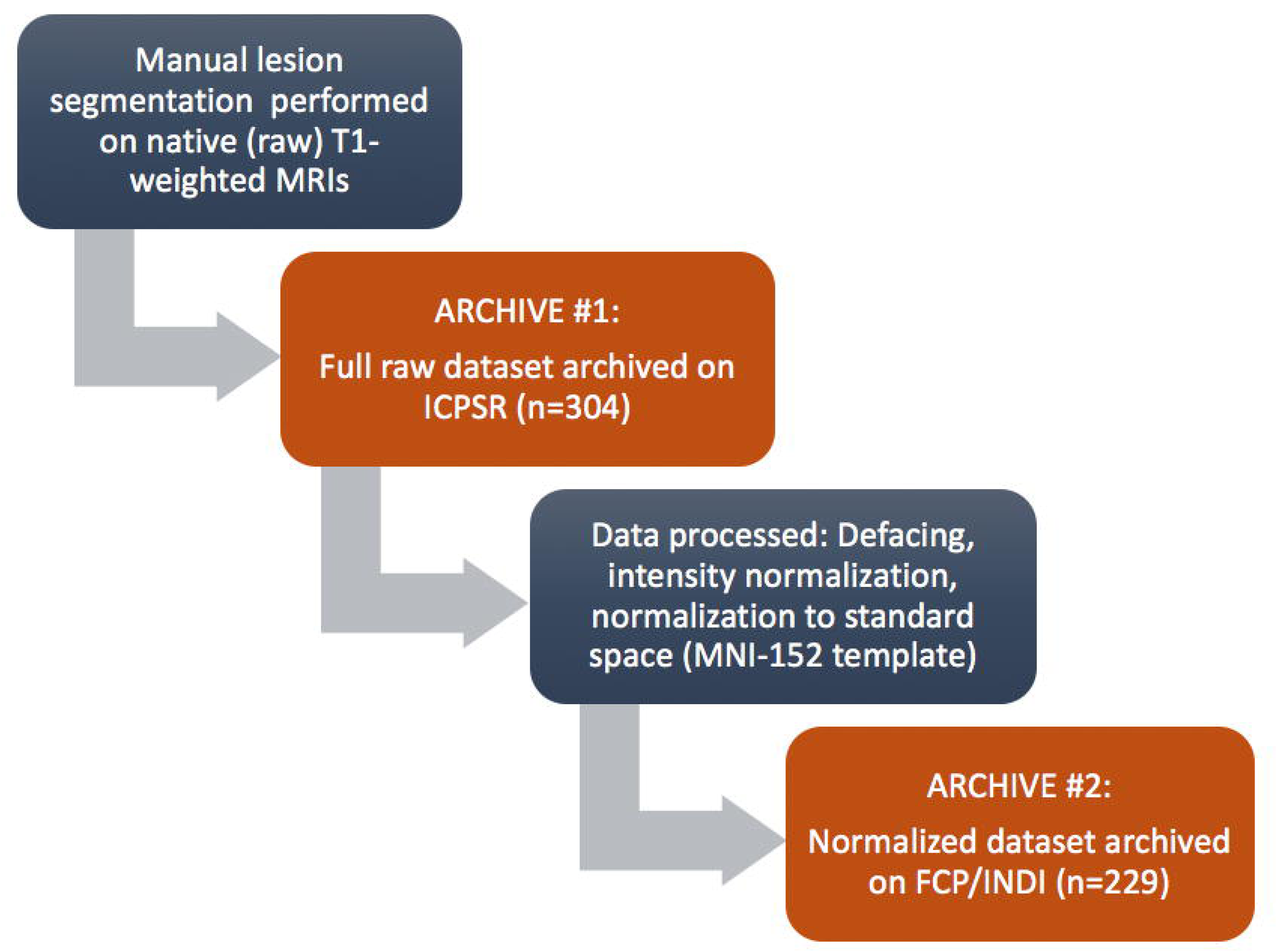
A schematic diagram showing the steps performed on the data for each archive release.

All ATLAS contributions were based on studies approved by local ethics committees and were conducted in accordance with the 1964 Declaration of Helsinki. Informed consent was obtained from all subjects. The receiving site’s ethics committee at the University of Southern California approved the receipt and sharing of the de-identified data. Data were fully de-identified by removing all 18 HIPAA (Health Insurance Portability and Accountability)-protected health information identifiers, lesions were manually segmented on each MRI, and all data were visually inspected before release. In addition, the terms of the data sharing agreements were approved by the University of Southern California’s technology transfer office.

### Data Characteristics

All T1-weighted MRI data were collected on 3T MRI scanners at a resolution of 1 mm^3^ (isotropic), with the exception of data from cohorts 1 and 2 which were collected on a 1.5T scanner with a resolution of 0.9 mm × 0.9 mm × 3.0 mm (excluded from the normalized dataset). Scanner information (scanner strength, brand) and image resolution are included in the ATLAS meta-data, and sample image header information for a subject from each of the cohorts can be found in *Supplementary Information*.

Characteristics of the ATLAS dataset include an average lesion volume across all cohorts of 2.128±3.898 × 10^4^ mm^3^, with a minimum lesion size of 10 mm^3^ and a maximum lesion size of 2.838 × 10^5^ mm^3^. Information regarding the distribution of lesions in the ATLAS dataset (e.g., single versus multiple lesions per individual, percent of lesions that are left versus right hemisphere, or subcortical versus cortical) can be found in Tables 1 and 2. Overall, slightly more than half of the subjects had only one lesion (58%) while the rest had multiple lesions (42.1%). Lesions were roughly equally distributed between left and right hemispheres (48.4% left hemisphere, 43.8% right hemisphere, 7.7% other location such as brainstem or cerebellum). In this dataset, there were more subcortical lesions than cortical lesions (70.7% subcortical, 21.5% cortical, 7.7% other).

**Table 1.** A total of 304 subjects within 11 cohorts were included in the full ATLAS Release 1.1 native dataset. The number of brains in which only one lesion was found (left/right hemispheres and other locations found within the brainstem and cerebellum, etc.), and the number of brains in which multiple lesions were found, are shown.

| Cohort | Number of Subjects | Brains with One Lesion |  |  | Brains with |
| --- | --- | --- | --- | --- | --- |
|  |  | Left | Right | Other |  |

|  |  |  |  |  | **Multiple Lesions** |
| --- | --- | --- | --- | --- | --- |
| c0001 | 6 | 3 | 2 | 0 | 1 |
| c0002 | 25 | 6 | 12 | 1 | 6 |
| c0003 | 55 | 14 | 19 | 0 | 22 |
| c0004 | 34 | 7 | 3 | 1 | 23 |
| c0005 | 30 | 9 | 3 | 6 | 12 |
| c0006 | 12 | 4 | 1 | 2 | 5 |
| c0007 | 36 | 9 | 9 | 0 | 18 |
| c0008 | 32 | 8 | 11 | 0 | 13 |
| c0009 | 12 | 8 | 0 | 0 | 4 |
| c0010 | 47 | 9 | 10 | 7 | 21 |
| c0011 | 15 | 1 | 11 | 0 | 3 |
| **Total** | **304** | **78** | **81** | **17** | **128** |
| **% Total Subjects** |  | **25.7%** | **26.7%** | **5.6%** | **42.1%** |

**Table 2.** The number of lesions found in each location (i.e. cortical vs. subcortical; left vs. right hemispheres), and other locations (i.e., brainstem, cerebellum, etc.) are shown. Here we have included primary lesions as well as additional lesions, resulting in 521 total lesion masks across n=304.

| **Cohort** | **Cortical Lesions** |  | **Subcortical Only Lesions** |  | **Other Lesion Locations** |
| --- | --- | --- | --- | --- | --- |
|  | **Left** | **Right** | **Left** | **Right** |  |
| c0001 | 2 | 2 | 2 | 1 | 0 |
| c0002 | 1 | 7 | 14 | 11 | 4 |
| c0003 | 3 | 0 | 38 | 44 | 0 |
| c0004 | 7 | 6 | 31 | 32 | 7 |
| c0005 | 9 | 1 | 20 | 14 | 7 |
| c0006 | 3 | 2 | 8 | 3 | 3 |
| c0007 | 8 | 10 | 28 | 18 | 4 |
| c0008 | 6 | 13 | 20 | 13 | 1 |
| c0009 | 5 | 3 | 12 | 2 | 0 |
| c0010 | 11 | 8 | 21 | 24 | 13 |
| c0011 | 0 | 5 | 3 | 10 | 1 |
| **Total** | **55** | **57** | **197** | **172** | **40** |
| **%Total Lesions** | **10.6%** | **10.9%** | **37.8%** | **32.9%** | **7.7%** |

### Training Individuals Performing Lesion Tracing

Eleven individuals were carefully trained in identifying and segmenting lesions. Individuals had a range of backgrounds, including undergraduate students, graduate students, and postdoctoral fellows. All tracers were given detailed information regarding neuroanatomy, and underwent standardized training, which utilized a detailed protocol as well as an instructional video. All tracers were guided through the training process with extensive feedback on lesion tracing performance by an expert tracer and in consultation with an expert neuroradiologist. The detailed protocol, with pictures of example tracings, is freely available and can be found on the ATLAS GitHub website (https://github.com/npnl/ATLAS/). All individuals were trained on an initial set of 5 brains with varying lesion sizes and locations (size range: min: 1,871 mm^3^, max: 162,015 mm^3^; location: cortical [Left: 0, Right: 1], subcortical [Left: 4, Right: 0]). After tracing the first set of 5 lesions, tracings were reviewed by an experienced tracer and differences in the lesion masks were discussed with the tracer. One week later, individuals retraced the lesions on the same set of 5 brains, but were blinded to their first segmentation attempt to examine intra-tracer reliability. After this, each lesion segmentation was reviewed by a separate tracer. In addition, the primary lesion location was identified by an expert neuroradiologist, who also created the meta-data (see *Metadata* below). Any questions regarding lesion masks were referred to the neuroradiologist. Inter- and intra-rater reliability measures and additional technical validation of the lesion tracings can be found in *Technical Validation* below. Finally, we note that lesion tracing is a subjective process, even across trained individuals. As mentioned in *Usage Notes* below, any problems, questions, or issues with specific lesion masks can be publically reported on our ATLAS GitHub under the Issues page (https://github.com/npnl/ATLAS/issues) so that the community of users can make comments and be aware of any identified issues. We will work to resolve any issues in a timely manner.

### Identifying and Tracing Lesions

To identify lesions, each T1-weighted MRI image was displayed using the multiple view option in MRIcron^20^, which displays the brain in the coronal, sagittal, and axial view (see Figure 2). To identify lesions, tracers looked for darker intensities within typically healthy tissue. For lesions that were more difficult to detect with the grayscale setting, colored look-up table settings (e.g., “cardiac”, “NIH”, or “spectrum” settings in MRIcron) were used to provide additional insight. Once the lesion or lesions were identified, the lesion mask was traced using either the coronal or axial view, using either a mouse, track pad, or tablet (i.e. Wacom Intuos Draw). A combination of MRIcron tools was used to draw the lesion masks, which included the 3D fill tool, the pen tool and the closed pen tool. Typically, and especially for larger sized lesions, tracers used the 3D fill tool to begin the segmentation. Crosshairs were placed in the center of the identified lesion and the tool would fill in voxels similar to the one at the point of origin with the selected radius and at the sensitivity specified by the difference from origin and difference at edge tools. The pen and closed pen tool, typically was used to fill in (or remove) the areas that the 3D tool had missed or was used to trace smaller lesions slice by slice. Once completed, lesion masks were saved in the volume of interest (VOI) file format with the identifier name “cXXXXsXXXXtXX_LesionRaw” (see *Data Records* and Table 3 below for full naming conventions). Lesions masks were then checked for correctness by a separate tracer, who made additional corrections to the lesion mask, if needed. After lesions were identified as being correct, masks were smoothed using MRIcron’s smooth VOI tool where the full width half maximum parameter was set to 2 mm and the threshold was set to 0.5. These masks were saved in both VOI and NIfTI file formats with the identifier name “cXXXXsXXXXtXX_LesionSmooth”.

**Table 3.** Filenames and file descriptions for ATLAS R1.1 dataset. * represents a wildcard.

**native folder (n=304)**
| Filename or Identifier | Description |
| --- | --- |
| cXXXXsXXXXtXX.nii.gz | Raw T1-weighted MRI for each subject, where c = cohort number, s = subject number, and t = time point |
| *LesionRaw.voi | Raw primary lesion mask, drawn as a volume of interest in MRICron |
| *LesionSmooth.voi | Smoothed primary lesion mask, drawn as a volume of interest in MRICron |
| *LesionSmooth.nii.gz | Smoothed primary lesion mask, saved as a nifti file |
| *LesionRaw/Smooth_1(or 2, 3, ...).voi/.nii.gz | Raw and smoothed secondary lesion masks (same as the three above, but for additional lesions) |

**standard folder (n=229)**
| Filename or Identifier | Description |
| --- | --- |
| Site, Subject ID, Session | Naming convention follows Brain Imaging Data Structure (BIDS) recommendations |

**Figure 2.**
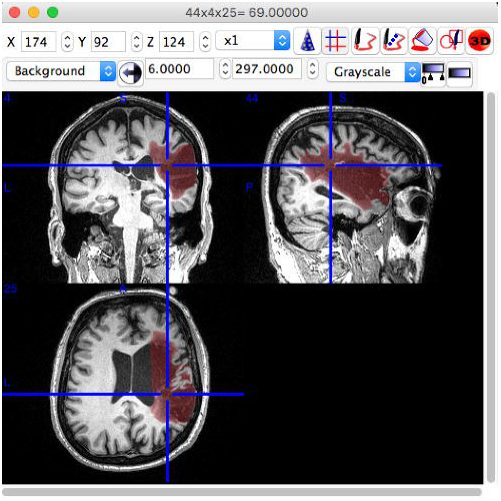
An example of lesion segmentation in MRICron.

Any additional lesions that were not contiguous with the primary lesion mask were drawn as separate lesion masks and labeled. As described in *Data Records*, any secondary lesions followed the same procedures as the primary lesion mask, but were labeled as Lesion_1, Lesion_2, Lesion_3 and so on, with the naming convention moving from the largest to smallest mask (e.g., Lesion_1 is the largest secondary lesion mask). In general, the primary lesion mask was the largest lesion, with any secondary lesion masks subsequently named and ordered by size (largest to smallest). The only exception to this was, in the case of multiple lesions, if the neuroradiologist identified a primary stroke location as a different lesion from the largest lesion mask. In these cases, we used the lesion identified by the neuroradiologist as the primary mask. This occurred in less than 5% of the subjects.

### Metadata

For each lesion, we also provided metadata on the lesion properties to give the user additional qualitative information, beyond the binary lesion mask. This information can be used to quickly sort the dataset based on specific lesion characteristics (e.g., only left hemisphere lesions, or only subcortical lesions). It can also provide additional insight into the types of lesions that succeed or fail for a given lesion segmentation algorithm. The lesion properties were manually reported for each individual lesion mask. These include the number of lesions identified and traced, and the location of each lesion (i.e. right/left, subcortical, cortical, or other). In order to count each lesion only once, we defined subcortical lesions as lesions that are contained completely in the white matter and subcortical structures. Any lesion that extends beyond this area and into the cortex is considered a cortical lesion. In this way, cortical lesions may extend into the subcortical space, but subcortical lesions do not extend into the cortical space. “Other” includes the brainstem and cerebellum. An experienced neuroradiologist also identified the following information for each individual brain: the type of stroke (e.g., embolic, hemorrhagic), primary stroke location, vascular territory, and intensity of white matter disease (periventricular hyperintensities, or PVH, and deep white matter hyperintensities, or DWMH). White matter hyperintensities were graded using the Fazekas scale^25^. For periventricular hyperintensities, the following grades were applied: 0 = absence, 1 = “caps” or pencil-thin lining, 2 = smooth “halo”, 3 = irregular PVH extending into the deep white matter. For deep white matter hyperintensities, the following grades were applied: a = absence, 1 = punctate foci, 2 = beginning confluence of foci, 3 = large confluent areas. The white matter hyperintensity ratings are included because areas of white matter hyperintensity often pose challenges for lesion segmentation algorithms. Finally, scanner strength, brand/model, and image resolution are included in the metadata as well.

*Normalization to a Standard Template, Intensity Normalization, and Defacing* To expand access to the dataset, we have also provided a subset of the data that is defaced, intensity-normalized, and normalized to standard (MNI-152) space. Lesion segmentation algorithms vary in whether the input should be in native (subject) space or a standardized space. Therefore, to provide this option for users, we also generated a version of the ATLAS dataset in standard space. To convert the images to standard space, MRI images first underwent automated correction for intensity non-uniformity and intensity standardization using custom scripts derived from the MINC-toolkit^26^ (https://github.com/BIC-MNI/minc-toolkit). These corrected images were linearly registered to the MNI-152 template using a version that was nonlinearly constructed and symmetric (version 2009; http://www.bic.mni.mcgill.ca/ServicesAtlases/ICBM152NLin2009) to normalize their intracranial volume in a standardized stereotaxic space^27^. Using the resulting transformation matrix, the labels drawn on the MRI images were also registered to the MNI template. The MRI images were resampled using the linear interpolation whereas their labels used a nearest neighborhood interpolation to keep their binary nature. Finally. Freesurfer’s mri_deface tools were used to perform the defacing (e.g., to remove any facial structures) (https://surfer.nmr.mgh.harvard.edu/fswiki/mri_deface) on all T1-weighted images.

Due to technical difficulties and differences in scanner image quality, a subset of brains is not included in the standard space conversion, resulting in a total of n=229 ATLAS brains converted into standard MNI space. Scans from the two cohorts with 0.9 × 0.9 × 3.0 mm resolution images, collected on 1.5T scanners, were excluded from this standardized dataset due to their lower resolution. In addition, any images that failed registration were excluded. Primary reasons for failed registration include large lesion volumes or poor image quality (e.g., image artifacts, motion artifacts). We are currently working on manually editing the registrations for these images, which will be released in the future. This dataset can be widely accessed from the FCP-INDI archive (see Figure 1 and Table 3 for archive details). All images were named in accordance with the INDI data policy, following the Brain Imaging Data Structure (BIDS), and a meta-data sheet using the INDI naming convention is included with this dataset.

### Probabilistic Spatial Mapping of ATLAS Lesion Labels

We also created a probabilistic spatial mapping of the lesion labels solely to visualize the distribution of lesion masks across the normalized ATLAS dataset. We note that this does not provide a representation of a true stroke distribution, but rather shows the distribution of lesions included in this dataset. To do this, we performed a population-based averaging of all the individual primary lesion labels in MNI space, producing a voxel-wise map where values can range from 0 at each voxel (always background for all subjects) to 1 (100% presence of the lesion label across subjects). A probabilistic spatial map of the primary lesions can be found in Figure 3 and a 3D visualization of the lesion map can be found in the following video link: https://www.youtube.com/watch?v=Ag5CUsRNY9Q. In addition, this map has also been provided in NIfTI format (.nii.gz) and uploaded to NeuroVault.org, an open-source database for neuroimaging data where it can be freely accessed (https://neurovault.org/collections/3073/).

**Figure 3.**
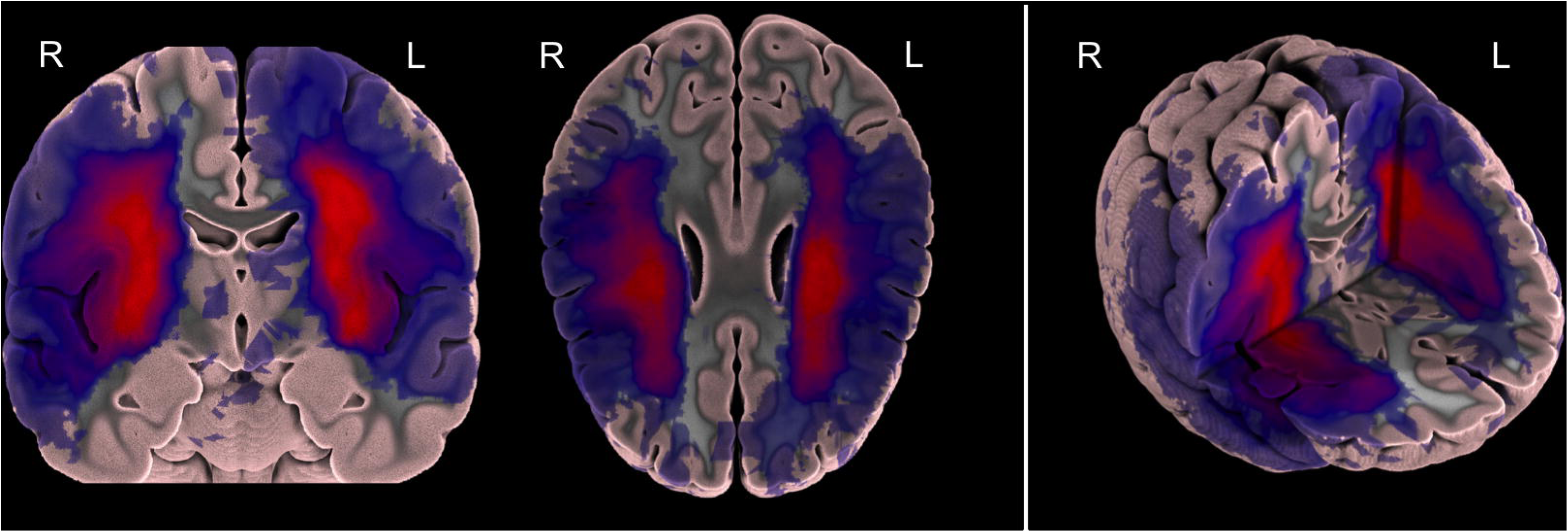
A probabilistic lesion overlap map for the primary lesions from the ATLAS R1.1 dataset. A 3D visualization of the lesion overlap map can be found at https://www.youtube.com/watch?v=Ag5CUsRNY9Q.

## Data Records

The full raw dataset (*native dataset*, n=304) is archived with the Archive of Disability Data to Enable Policy research at the Inter-university Consortium for Political and Social Research (ICPSR). ICPSR is the world’s largest social science data archive that supports several substantive-area archive collections including disability and rehabilitation. ICPSR provides access to the data and provides technical assistance to individuals accessing the data.

In addition, a standardized, defaced subset of the dataset (*standardized* dataset, n=229) is archived with the International Data Sharing Initiative, which hosts many widely available neuroimaging datasets such as the Functional Connectome Project (FCP-INDI). See *Usage Notes* for more details regarding access.

For the full dataset archived with ICPSR, the naming convention and description of the files in ATLAS R1.1 can be found in Table 3. Within the ATLAS R1.1 main folder, there is an excel file with the metadata for the entire dataset. The data in this archive is in native space (i.e., original subject space; n=304). Throughout the dataset, MRIs are named and sorted based on each cohort (c); each cohort is in the format of cXXXX where XXXX is the number that the cohort was assigned (e.g., c0001). There are 11 total cohorts. Within each cohort folder are the individual subject(s) folders. Subject folders are named based on the cohort that they are in (cXXXX), the subject number that they were assigned (sXXXX) and the time point at which they were taken (tXX) (e.g. c0001s0004t01). For instance, participants with data taken two weeks apart would have two time points, where t01 is the first time point and t02 is the second. Every image starts with the subject identifier of cXXXXsXXXXtXX.

Each subject folder has several components: at minimum, each will have the original T1-weighted MRI image (*.nii.gz) and three masks for the main lesion: the unsmoothed lesion mask (*LesionRaw.voi), and two smoothed lesion masks in .voi and .nii.gz formats (*LesionSmooth.voi; *LesionSmooth.nii.gz). The LesionRaw volume is the original hand-traced lesion volume, while the LesionSmooth volume used a Gaussian smoothing kernel (full width half maximum parameter set to 2 mm, threshold set to 0.5, to overcome small errors between slices in tracing; see *Methods* above). We anticipated that most researchers would use the LesionSmooth volume as it is slightly more robust to small slice-by-slice human errors, and therefore created the .nii.giz version from this. Notably, the .voi files are in an MRIcron format so the masks can be further edited in MRICron if desired. The .nii.gz files use the standard NIfTI format^28^ (http://nifti.nimh.nih.gov/nifti-1/), which can be opened, edited, and viewed by most standard neuroimaging software.

If a particular subject had multiple lesions, for each additional lesion, there would be three additional lesion masks (e.g. *LesionRaw_1.voi, *LesionSmooth_1.voi, *LesionSmooth_1.nii.gz). In general, lesions were ranked based on size where the largest lesion was considered the main lesion. As mentioned previously, if the largest lesion differed from the primary lesion identified by the neuroradiologist, we deemed the primary lesion to be the one identified by the neuroradiologist. This occurred in less than 5% of cases.

Finally, in the FCP-INDI archive (standardized dataset, n=229), there is a separate naming convention, following the Brain Imaging Data Structure (http://bids.neuroimaging.io/), adopted by FCP-INDI. Images in this dataset have been normalized to a standard MNI-152 template, intensity normalized, and defaced. Table 3 provides a list of all naming conventions and filenames, along with descriptions.

## Technical Validation

Each trained tracer created lesion masks for the same five brains twice, one week apart, to assess both inter- and intra-tracer reliability. Training lesions ranged in size and difficulty (see *Methods*). Each tracer’s lesion masks were compared, providing both inter- and intra-rater reproducibility measures. We first calculated inter- and intra-rater reliability measures using the lesion volumes. Based on lesion volumes, the inter-rater reliability was 0.76±0.14, while the intra-rater reliability was 0.84±0.09.

In addition, we also calculated inter-and intra-rater reliability using the Dice similarity coefficient (DC), which is a segmentation accuracy metric, and Hausdorff’s distance (HD), which is a metric of the maximum distance between two volumes surface points. DC allows us to examine not only if the volumes are similar, but also if the same voxels are being selected as part of the lesion mask or not. This is particularly useful for comparing neuroimaging volumes, such as lesion masks. DC is calculated by the formula:

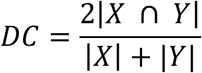

where X and Y represent the voxels from each lesion segmentation, and DC ranges from 0 to 1 (where 0 means there were no overlapping voxels and 1 means that the segmentations were completely the same). HD allows us to examine the distance between the surfaces of two images and thus can be used to identify outliers, providing another useful metric for comparing neuroimaging volumes. HD is calculated by the formula:

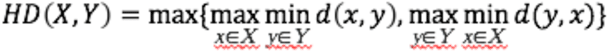

where x and y represent the surface points from the volumes X and Y respectively. HD is measured in millimeters, and a lower value denotes that the maximum distance between the two images is smaller.

Inter-rater scores (DC, HD) were calculated for each manual segmentation by comparing each individual tracer’s lesion mask to the rest of the tracers’ lesion masks. Inter-rater DC and inter-rater HD scores were then averaged to obtain one final DC score and one final HD score for the initial segmentations (average inter-rater DC for first segmentation: 0.73±0.20; average inter-rater HD for first segmentation: 22.57±21.36mm) and for the second segmentations (average inter-rater DC for second segmentation: average inter-rater HD for second segmentations: 25.29±23.53mm). Furthermore, intra-rater DC and HD scores were calculated for each brain traced by comparing the initial segmentation to the secondary segmentation for each tracer; these scores were then averaged to obtain a final intra-rater DC score (0.83±0.13) and a final intra-rater HD value (21.02±22.66mm).

Trained tracers segmented all lesion masks. In addition, each lesion mask was checked by a separate tracer, and changes were made to the lesion mask as needed. Any difficulty identifying the lesion was discussed with the expert neuroradiologist. Lastly, after the completion of the dataset, lesion masks were checked a second time to ensure correct segmentation and data descriptors. It is important to note that while tracers did participate in a thorough training process and segmentations were checked multiple times, this is still a subjective process. Comments regarding the lesion masks can be submitted as issues on the ATLAS GitHub site (https://github.com/npnl/ATLAS/issues), and we plan to publish updated and expanded versions of this dataset based on feedback and comments from users (see *Usage Notes*).

## Usage Notes

The full native-space archived dataset (n=304) can be found at ICPSR: http://www.icpsr.umich.edu/icpsrweb/ICPSR/studies/36684. For more information on the data archive, visit the ICPSR website (https://www.icpsr.umich.edu/icpsrweb).

In addition, a standardized, intensity-normalized, defaced subset of the data (n=229) can be found at FCP-INDI: http://fcon_1000.projects.nitrc.org/indi/retro/atlas.html.

Data is accessible under a standard Data Use Agreement, under which users must agree to only use the data for purposes as described in the agreement. Users of the ATLAS dataset should acknowledge the contributions of the original authors and research labs by properly citing this article and the data repository link from which they accessed the data.

As described above, lesions were segmented using the NITRC open source software MRIcron which can be downloaded from the NITRC website (https://www.nitrc.org/projects/mricron). Users can also quickly and easily view the brains on BrainBox (http://brainbox.pasteur.fr/), an open-source Web application to collaboratively annotate and segment neuroimaging data available online^29^. For additional quick quantification, our group has also created a small package of scripts called SRQL (Semi-automated Robust Quantification of Lesions), which provide three features: it uses a semi-automated white matter intensity correction to further correct for human errors in lesion tracing, outputs a report of descriptive statistics on lesions (hemisphere and volume of lesion), and gives users the option to perform analyses in native or standard space (https://github.com/npnl/SRQL)^30^. In addition, as we plan to grow this dataset in the future, additional releases of data or software will be announced on our ATLAS GitHub page (https://github.com/npnl/ATLAS/). Any issues or feedback can also be submitted on the ATLAS GitHub page under “issues,” and a team of researchers will address these in a timely manner. Finally, as a general note regarding the usage of this dataset, we strongly encourage users to be cautious of overfitting training algorithms to this particular dataset. We note that this data is relatively diverse, given the data collection across 11 research sites worldwide. However, we caution users against overfitting to only a particular cohort or subset of this data. Future work will aim to provide additional test datasets for users to properly test their algorithms on untrained data.

## Acknowledgments

We thank Dr. Mauricio Reyes for insightful conversations and would like to acknowledge the following people for their assistance on this effort: Anthony Benitez, Xiaoyu Chen, Cristi Magracia, Ryan Mori, Dhanashree Potdar, Sandyha Prathap. The archiving of this dataset was specifically supported by the NIH-funded Center for Large Data Research and Data Sharing in Rehabilitation (CLDR; https://www.utmb.edu/cldr) under a Category 2 Pilot Grant (P2CHD06570) and this work was also funded by an NIH K01 award (K01HD091283).

## Author contributions

S.-L.L. conceptualized the study, reviewed lesions, analyzed data, established archives, and contributed to the writing and editing of the manuscript. J.M.A. segmented and reviewed lesions, oversaw the organization to the segmentation process and contributed to the writing and editing of the manuscript. N.W.B. organized, segmented and reviewed lesions. M.S. provided the neuroradiology expertise and information. K.L.I. and H.K. performed data analysis. H.K. also performed data processing and generated the standardized dataset and probabilistic lesion maps. T.A. provided data visualization expertise and generated the figures/videos. J.C., D.S., A.S. J.I., C.J., W.N., D.V. and S.L. segmented and/or reviewed lesions. P.H., B.K., N.K., L.A.-Z., S.C.C., J.L., S.S., L.T.W., J.W., C.W., C.Y. collected and provided the MRI data. M.L., A.P., and A.S. handled the archiving of the data.

## References

1. Benjamin, E.J., et al. Heart Disease and Stroke Statistics-2017 Update: A Report From the American Heart Association. Circulation 135, e146–e603 (2017).

2. Feigin, V.L., et al. Global and regional burden of stroke during 1990-2010: findings from the Global Burden of Disease Study 2010. Lancet (London, England) 383, 245–254 (2014).

3. Kwakkel, G., Kollen, B.J., Van der Grond, J.V. & Prevo, A.J.H. Probability of regaining dexterity in the flaccid upper limb: Impact of severity of paresis and time since onset in acute stroke. Stroke 34, 2181–2186 (2003).

4. Ren, J., Kaplan, P.L., Charette, M.F., Speller, H. & Finklestein, S.P. Time window of intracisternal osteogenic protein-1 in enhancing functional recovery after stroke. Neuropharmacology 39, 860–865 (2000).

5. Group, I.-C. Association between brain imaging signs, early and late outcomes, and response to intravenous alteplase after acute ischaemic stroke in the third International Stroke Trial (IST-3): secondary analysis of a randomised controlled trial. The Lancet Neurology 14, 485–496 (2015).

6. Burke Quinlan, E., et al. Neural function, injury, and stroke subtype predict treatment gains after stroke. Ann Neurol 77, 132–145 (2015).

7. Marie-Héléne, M. & Cramer, S.C. Biomarkers of recovery after stroke. Current opinion in neurology 21, 654–659 (2008).

8. Nijland, R.H.M., van Wegen, E.E.H., Harmeling-van der Wel, B.C. & Kwakkel, G. Presence of finger extension and shoulder abduction within 72 Hours after stroke predicts functional recovery. Stroke 41, 745–750 (2010).

9. Riley, J.D., et al. Anatomy of stroke injury predicts gains from therapy. Stroke 42, 421–426 (2011).

10. Cramer, S.C., et al. Predicting functional gains in a stroke trial. Stroke 38, 2108–2114 (2007).

11. Jongbloed, L.Y.N. Prediction of function after stroke: a critical review. Stroke 17, 765–776 (1986).

12. Nouri, S. & Cramer, S.C. Anatomy and physiology predict response to motor cortex stimulation after stroke. Neurology 77, 1076–1083 (2011).

13. Prabhakaran, S., et al. Inter-individual variability in the capacity for motor recovery after ischemic stroke. Neurorehabilitation and Neural Repair 22, 64–71 (2007).

14. Stinear, C. Prediction of recovery of motor function after stroke. The Lancet Neurology 9, 1228–1232 (2010).

15. Zhu, L.L., Lindenberg, R., Alexander, M.P. & Schlaug, G. Lesion load of the corticospinal tract predicts motor impairment in chronic stroke. Stroke 41, 910–915 (2010).

16. Fiez, J.A., Damasio, H. & Grabowski, T.J. Lesion segmentation and manual warping to a reference brain: Intra- and interobserver reliability. Human Brain Mapping 9, 192–211 (2000).

17. Montaner, J., et al. Plasmatic level of neuroinflammatory markers predict the extent of diffusion-weighted image lesions in hyperacute stroke. Journal of Cerebral Blood Flow & Metabolism 23, 1403–1407 (2003).

18. Sakamoto, Y., et al. Early ischaemic diffusion lesion reduction in patients treated with intravenous tissue plasminogen activator: infrequent, but significantly associated with recanalization. International Journal of Stroke 8, 321–326 (2013).

19. Thomas, R.G.R., et al. Apparent diffusion coefficient thresholds and diffusion lesion volume in acute stroke. Journal of Stroke and Cerebrovascular Diseases 22, 906–909 (2013).

20. Wittsack, H.-J., et al. MR Imaging in Acute Stroke: Diffusion-weighted and Perfusion Imaging Parameters for Predicting Infarct Size. Radiology 222, 397–403 (2002).

21. Pustina, D., et al. Automated segmentation of chronic stroke lesions using LINDA: Lesion identification with neighborhood data analysis. Human Brain Mapping 37, 1405–1421 (2016).

22. de Haan, B., Clas, P., Juenger, H., Wilke, M. & Karnath, H.-O. Fast semi-automated lesion demarcation in stroke. NeuroImage. Clinical 9, 69–74 (2015).

23. Maier, O., et al. ISLES 2015 - A public evaluation benchmark for ischemic stroke lesion segmentation from multispectral MRI. Medical Image Analysis 35, 250–269 (2017).

24. Rorden, C. & Brett, M. Stereotaxic display of brain lesions. Behav Neurol 12, 191–200 (2000).

25. Fazekas, F., Chawluk, J., Alavi, A., Hurtig, H. & Zimmerman, R. MR signal abnormalities at 1.5 T in Alzheimer's dementia and normal aging. American Journal of Roentgenology 149, 351–356 (1987).

26. Sled, J.G., Zijdenbos, A.P. & Evans, A.C. A nonparametric method for automatic correction of intensity nonuniformity in MRI data. IEEE Transactions on Medical Imaging 17, 87–97 (1998).

27. Collins, L., D., Neelin, P., Peters, T.. & Evans, A., C. Automatic 3D intersubject registration of MR volumetric data in standardized Talairach space. Journal of Computer Assisted Tomography 18, 192–205 (1994).

28. Cox, R.W., et al. A (sort of) new image data format standard: Nifti-1. Neuroimage 22, e1440 (2004).

29. Heuer, K., Ghosh, S., Robinson Sterling, A. & Toro, R. Open Neuroimaging Laboratory. Research Ideas and Outcomes 2, e9113 (2016).

30. Ito, K., Anglin, J. & Liew, S.-L. Semi-automated Robust Quantification of Lesions (SRQL) Toolbox. Research Ideas and Outcomes 3, e12259 (2017).

